# Beyond endemism, expanding conservation efforts: a relictual Pleistocene distribution and first report of the prickly pear cactus, *Opuntia bonaerensis,* in Brazil and Uruguay

**DOI:** 10.1101/2020.03.06.981480

**Authors:** Matias Köhler, Luíz F. Esser, Fabián Font, Tatiana T. Souza-Chies, Lucas C. Majure

## Abstract

Geographical range is one of the critical features for species conservation assessment. Nevertheless, species distribution is frequently unknown, undervalued or overlooked. During a broad taxonomic and floristic study of the southern South American prickly pear species (*Opuntia* spp.), new records of a previously endemic Argentinian taxon have been found in Uruguay and Brazil. Molecular phylogenetic inference was carried out to further evaluate the identity of the new records, and ecological niche models were implemented to test how the new records would fit in the previous known distribution of the species. Through molecular and morphological evidence, we confirmed the new records of *Opuntia bonaerensis* for Brazilian and Uruguayan floras and discussed its phylogenetic relationship and morphologic similarities with closely related species. Our new records uncovered a distributional pattern congruent with the Neotropical Peripampasic Orogenic Arc, which must be further explored to better determine the biogeographic history of the species. Ecological niche models (ENM) revealed that *O. bonaerensis* had a putative ancient distribution across the grasslands and shrublands in the Pampean region largely congruent with the populations found in Brazil and Uruguay, suggesting relictual Pleistocene populations of the species and the role of glacial/interglacial cycles on the distribution of the species. In a prospective climate change scenario, ENM suggests that the species would in general be more restricted to the southernmost limits of the Pampa region and previous outlying records from Mendoza (Argentina) are a putative future refuge for *O. bonaerensis*. The importance of these new records for biodiversity and conservation assessment efforts that are ongoing at different scales in Brazil and neighbor areas is highlighted.

## Introduction

Cacti are a conspicuous and diverse family of Angiosperms composed of roughly 1500 species distributed throughout the entire American continent (Britton and Rose, 1919; Anderson, 2001; Hunt et al., 2006; Guerrero et al., 2018). The clade is remarkable in showing intriguing modifications to survive in extremely adverse environments with drought and arid conditions usually exhibiting succulence accompanied by spines (Mauseth, 2006), but it also occurs in tropical and wet environments, especially as epiphytes (Anderson, 2001; Taylor and Zappi, 2004). The cactus lineage diverged from its closest relatives around 35 million years ago (Mya) (Arakaki et al., 2011; Hernández-Hernández et al., 2014). However, there is a time lag between the origin and extant diversification of Cactaceae, with the latter taking place mainly during the last 10 Mya, in which across disparate clades, the family has experienced high and differential rates of diversification (Arakaki et al., 2011; Hernández-Hernández et al., 2014). Impacts of the Pleistocene glacial/interglacial cycles have also been documented driving the diversification and distribution of some groups (Majure et al., 2012a, 2012b; Ornelas and Rodríguez-Gómez, 2015; Franco et al., 2017b; Silva et al., 2018a).

Evaluation of the conservation status of cacti has recently captured much attention, revealing the group as the fifth most endangered clade of any major taxonomic group, with 31% of evaluated species under some categories of threat (Goettsch et al., 2015). Several threats are driven by human activities accompanying land conversion to agriculture, unscrupulous collection of live plants as biological resources for the horticultural trade and private ornamental collections, and residential and commercial development (Oldfieldb, 1997; Goettsch et al., 2015). However, although cacti are an iconic group and broadly call attention from scientists and cactus aficionados, reliable information regarding species limits and their geographic distribution throughout the family are frequently unavailable or deficient (Zappi et al., 2011). In fact, succulence, and especially spines, have made cacti an intimidating group for botanists to collect, resulting in herbaria with very deficient representation of cactus specimens or very poorly prepared specimens with many gaps to be filled (Reyes-Agüero et al., 2007; Hunt, 2014; Majure et al., 2017; Zappi et al., 2018).

*Opuntia* Mill. s.str. is the second most speciose genus of the family (after *Mammillaria* Haw.), containing around 180 species, with a broad distribution across all the Americas from Argentina to Canada, including Central America and Caribbean region (Anderson, 2001; Hunt et al., 2006; Majure et al., 2012a). The group has a putative origin on southern South America and subsequently dispersal events of lineages to Northern South America, the Caribbean region, Central America and to the North American deserts (Majure et al., 2012a). Members of *Opuntia* share a combination of morphological traits, including sympodial shrubs or trees with flattened photosynthetic stems (cladodes), areoles with smooth or retrorsely barbed spines and glochids (small, hair-like spines), reduced and ephemeral leaves, radial, diurnal flowers with inferior ovaries and multilobed stigmas, stamens frequently thigmonastic, reticulate semitectate pollen, and seeds covered by a sclerified, funicular aril (Buxbaum, 1953; Anderson, 2001; Stuppy, 2002; Hunt et al., 2006; Majure and Puente, 2014; Majure *et al.,* 2017). Eight major clades are recognized within *Opuntia* (Köhler et al., *in prep.*), which exhibit a variety of morphological characters such as dioecy, humming bird pollination, dry fruit, epidermal and seed pubescence, as well as rhizomes and tuberous roots that are unique to some species in the genus (Britton and Rose, 1919; Majure et al., 2012a; Majure and Puente 2014; Majure et al., 2017).

Conservation status assessments of the prickly pear cacti (*Opuntia* spp.) are relatively incipient (Goettsch et al., 2015) and are not a simple issue. Geographical range is a critical feature for conservation assessment of the International Union for the Conservation of Nature - IUCN (Criteria B and D2, IUCN 2001), and it has been the most used criterion to assess threatened plant species in one of the categories of extinction risk (Collen et al., 2016). However, many *Opuntia* species are worldwide cultivated for different purposes (such as fruit and vegetable crops, forage and fodder for livestock and ornamentals (Inglese et al., 2002; Nefzaoui and Salem, 2002)), increasing the complexity of knowledge regarding the distribution of some species. So, the lack of data regarding the distribution of species has an immense impact on those evaluations. There are currently five of the 84 evaluated species under threat in some of the three criteria of extinction risk (EN, VU, CR) of the IUCN, and ten taxa are data deficient (DD) (IUCN 2019), and many taxa remain unevaluated. Morphologically variable species, and frequent hybridization make species delimitation within *Opuntia* a problematic issue that also is reflected in their conservation and biodiversity management.

Ecological niche models (ENM), which are produced by combining species occurrence data with environmental data layers, have transformed evolutionary, systematics and conservation biodiversity studies across disparate organisms (Raxworthy et al., 2007, Peterson 2001, Kozak et al., 2008). ENM have allowed scientists to develop more reliable hypotheses to describe, understand and predict geographic and environmental distributions of species to the present, as well as to the past and future scenarios (Peterson 2006), becoming a powerful tool to infer local adaptations (Rolland et al., 2015), environmental drivers of diversity (Barros et al., 2015), interglacial microrefugia in paleoenvironments (Bonatelli et al., 2014) and impacts of climate change on species distribution (Maguire et al., 2015). Besides being descriptive, novel approaches have been used ENM to quantitatively test niche differences, such as niche overlap, niche equivalency and niche similarity (Warren et al., 2008; Broennimann et al., 2012; Dagnino et al., 2017), increasing the significance of niche modelling applications.

*Opuntia bonaerensis* Speg. is a species described for the Argentinean flora in the early 20th century (Spegazzini, 1901). When described, it was mentioned to be rare, since it was observed at only a few localities in the southern Buenos Aires province (Sierras of Curamalál and Tornquist). The taxon has a complicated taxonomic history being circumscribed in various ways in seemingly arbitrary taxonomic treatments (Spegazzini, 1905; Britton and Rose, 1919; Spegazzini, 1925; Leuenberger, 2002), which is not uncommon in the taxonomic history of *Opuntia* (Hunt, 2002). Just recently, in the taxonomic revision of the *Opuntia* series Elatae, *O. bonaerensis* was resurrected as an endemic species of the Argentinian pampean region and delimited using a set of diagnostic morphological characters that have been historically ignored for southern South American species, such as bud flower apices, stigma colors and color of the inner pericarpel tissue (Font 2014; Las Peñas et al., 2017). The species was further included in a preliminary phylogenetic study of the southern South American species of *Opuntia*, confirming it assignment to this taxonomic series (Realini et al. 2014b).

During a broad floristic and taxonomic study of *Opuntia* in southern South America, unexpected populations of *O. bonaerensis* have been found outside of its previously known Argentinian distribution, in the Uruguayan and Brazilian pampean regions. Considering the general difficulty in identifying *Opuntia* species using morphological characters, we further tested the lineage identity of the groups under study using molecular phylogenetic inference of populations across the range of the taxon. To test how these unexpected records outside of Argentina could be explained in the natural history of the species, we generated four datasets of different distribution records in which we explored ENM with projections to past, current and future climate scenarios. Confirming the new records of *Opuntia bonaerensis* in Uruguay and Brazil using both morphological characters and molecular phylogenetics, as well as a putative relictual Pleistocene distribution in these regions, we highlight implications for conservation efforts in the regional floras of Brazil and neighboring areas considering the current global and local strategies for biodiversity conservation.

## Material and methods

### Studied area and data collection

Extensive fieldworks were carried out in southern South America encompassing the main natural ecoregions to obtain data about natural populations of *Opuntia*. The region is represented by subtropical grasslands permeated by rocky outcrops that compose the Pampa biome or Río de la Plata grassland (Andrade et al., 2018) and the Chaco region (Pennington et al., 2000). The major herbaria from the region have been examined to check distribution records and specimen identification of *Opuntia,* including unidentified materials: BA, BAF, BCWL, CORD, CTES, HAS, ICN, LIL, LP, MBM, MCN, MVFA, MVJB, MVM, SI (acronyms according Thiers 2019+, except BCWL, non-indexed herbarium of the Biological Control of Weeds Laboratory (FuEDEI), Hurlingham, Buenos Aires, Argentina). The digital database of Brazilian collections was also consulted through the SpeciesLink platform (2019) to check herbaria from disparate geographical regions.

Herbarium materials were examined to obtain data regarding morphological features as well as other important information, such as ecological, biological and distributional data (Vogel, 1987). Likewise, fieldwork was carried out to obtain morphological, as well as biological (e.g., phenology, pollinators) and ecological (e.g., soil and vegetation type) data across populations.

Samples of materials collected in the field were dried in silica gel to keep tissues available for further molecular studies (Funk et al., 2017), and representative materials were collected as voucher (see Table S2 for further information). Morphological characters were assessed based on commonly used characters for prickly pears identification (e.g., cladodes morphology, habit and growth form, spine production) (Pinkava, 2003, Majure et al. 2017), with a special focus on those reported by Font (2014) for the southern South American species, such as bud flower apices, stigma color, and inner pericarpel tissue color. The criteria for identification of southern South American species of *Opuntia* followed those proposed by Font (2014) and Las Peñas et al., (2017).

### DNA sampling, sequence alignment and phylogenetic analyses

A small dataset was selected to test the hypothesis of whether the new Uruguayan and Brazilian records represented lineages of typical *Opuntia bonaerensis* from Argentina. One sample of each of those new records was selected to sequencing, and one sample of *O. bonaerensis* from the type locality in Argentina was also incorporated. We included a representative dataset of all southern South American species from *Elatae* series (sensu Font, 2014) to contextualize the phylogenetic relationships between the taxa sampled. Additionally, two North American taxa *(Opuntia macrorhiza* Engelm. and *Opuntia austrina* Small) and a sample of *Brasiliopuntia brasiliensis* (Willd.) A. Berger were selected as outgroups according to previous phylogenetic studies in the group (Majure et al., 2012a; Majure and Puente 2014). The complete information regarding the taxa sampled is presented in the supplemental Table S1.

Cladode epidermal tissue of selected taxa was used for DNA extraction using a standard CTAB incubation (Doyle and Doyle, 1987) followed by chloroform/isoamyl alcohol precipitation and silica column-based purification steps as described in Neubig et al. (2014) and Majure et al. (2019). Whole genomic DNAs were quantified using the Qubit dsDNA BR Assay Kit and Qubit 2.0 Fluorometer (Life Technologies, Carlsbad, California, USA); high-molecular-weight DNA (>15 kb) samples showing no degradation were considered suitable and then sent to Rapid Genomics LLC (http://rapidgenomics.com/home/; Gainesville, FL.) for library preparation and high-throughput sequencing using the Illumina HiSeq X platform with 150 bp paired-end reads.

Raw reads were imported into Geneious 11.1.5 (Biomatters, Auckland, New Zealand) and set paired reads with an expected insert size of 300 bp calculated with BBMap (Bushnell 2016). Low quality bases (Q & 20) were trimmed and all reads shorter than 20 bp were discarded using BBDuk for quality control of the reads (Bushnell 2016). Then, a reference guided assembly was carried out on the trimmed reads using selected phylogenetically informative regions from the chloroplast genome as a reference (see below). Based on previous molecular phylogenetic studies in *Opuntia* and related cacti (Arakaki et al., 2011, Hernández-Hernández et al., 2011, Majure et al., 2012a, Hernández-Hernández et al., 2014), we selected the chloroplast genes *ccsA*, *rpl16, trnK* including *matK* (*trnK/matK*) and the intergenic spacer *trnL-trnF* as markers, using those GeneBank sequences as references. The reference mapping pipeline was conducted using the Geneious mapper feature with a medium-low sensitivity, and we generated a majority consensus sequence from the reference-mapped raw reads.

Sequences of each marker from each taxon were concatenated as one sequence, and a multiple sequence alignment was performed across all samples using the MAFFT v. 7 (Katoh and Standley 2016) plugin in Geneious with default settings, and then manually corrected. Phylogenetic inference was performed using the Maximum Likelihood (ML) approach implemented in RAxML 8.2.4 (Stamatakis 2014) on the CIPRES Science Gateway Web Portal (Miller et al., 2010) using the GTR+G model of molecular evolution, treating all gaps as missing data. Support values were estimated undertaking 1,000 bootstrap replicates.

### Building, assembling, projecting and exploring ecological niche models

As presence records, four approaches were considered, and are summarized in Fig. 1: (1) using data previously known as the natural distribution of the species, gathered from our herbarium studies, but segregating records from Mendoza (Argentina), which could be environmental outliers from non-natural distribution (see Font, 2014) (henceforth, **P** data, 17 records); (2) summing data previously known (**P**) and new records from our field expeditions in Uruguay and Brazil (henceforth, **PN** data, 25 records); (3) as an alternate approach, we also clustered Mendoza and previously known records (henceforth, **PM** data, 19 records); and (4) the sum of previously known, new data and Mendoza records (henceforth, **PMN** data, 27 records). This segregation was important to track environmental contribution from each set, as well as to determinate whether they are more different than expected by chance.

**Figure 1.**
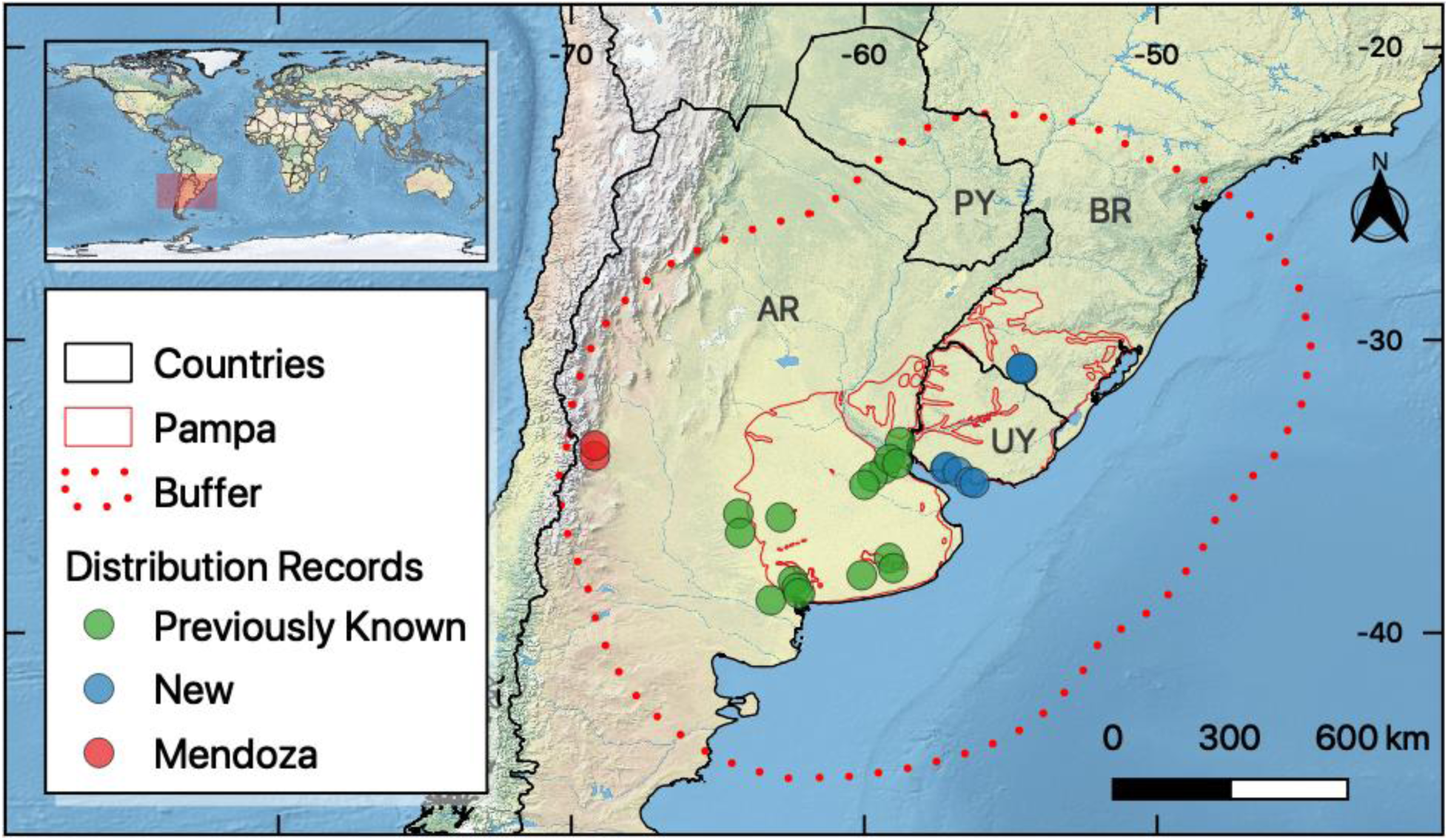
Distribution records of *Opuntia bonaerensis* (Cactaceae). Green circles represent the previously known distribution of the species in Argentina (AR), while blue circles in Uruguay (UY) and Brazil (BR) are newly reported here. Red circles represent previously known records from Mendoza, which are environmentally distinct populations. (For interpretation of the references to color in this figure legend, the reader is referred to the web version of this article).

Considering that little is known about the biology of *Opuntia bonaerensis* and its relationship with bioclimatic variables, we used PCA-axes as a proxy of ecological niche models, as they summarize species relationship with environment variables. We generated a correlation matrix of standardized variables, which was composed using all bioclimatic variables from WorldClim v.1.4 (30” resolution, Hijmans et al. 2005). In total, three axes were selected, which accounted for 89% of the overall variation. We could not use more than three, since the use of more variables could result in overparameterization. Axes were projected to the past, current and future scenarios using ENMGadgets package (Barve and Barve, 2016) on R (R Core Team, 2017). Past scenarios comprised the Last Glacial Maximum (LGM: about 21,000 years ago - 21 kya) and Mid-Holocene (MID: about 6 kya), while future scenarios comprised the extreme scenarios RCP 2.6 and RCP 8.5 to the year of 2070 (IPCC 2013). The extreme scenarios were selected to allow us to infer the putative minimal and maximal impacts of climate change on the species distribution. To keep comparability between scenarios and reduce uncertainties we used two General Circulation Models (CCSM4 and MIROC-ESM) from WorldClim v.1.4 (2.5 arc-minutes resolution, Hijmans et al. 2005), which were the only two available to every scenario calculated.

Modeling domain comprised a 6-degree wide buffer in the region of the Río de La Plata grasslands/Pampa (Dixon et al. 2014; Andrade et al. 2018), which is the previously known distribution of the species (Font, 2014), with a buffer that comprises Mendoza records. Models were generated using ten algorithms available in *biomod2* R package, to be known: Artificial Neural Network, ANN, Classification Tree Analysis, CTA, Flexible Discriminant Analysis, FDA, Generalized Additive Model, GAM, Generalized Boosting Model, GBM, Generalized Linear Model, GLM, Surface Range Envelope, SRE, Multiple Adaptive Regression Splines, MARS, Random Forest, RF, and MaxEnt (Thuiller et al., 2016). Pseudoabsence selection was performed in groups following Barbet-Massin et al. (2012). The first group comprised GLM, GAM and MaxEnt, where we selected 1000 pseudoabsences disposed randomly in environmental space. Afterwards, we generated a two-degree wide buffer from each presence point and selected pseudoabsences for the other two groups, as follows: for the second group, which comprised FDA, MARS and ANN, we selected 100 pseudoabsences, while for the third group (CTA, BRT, RF and SRE) we selected the same number of presence records. This routine was applied to every dataset separately, as they differed in number and location of presence records. We made 10 independent runs of 4-fold cross-validation, keeping 75% of data to build the models and 25% to test them, summing a total of 400 models for each dataset. Ensemble models were built through a committee average approach, where projections are binarized using a TSS threshold and summed (see further cutoff values in Tab. 1), resulting in a map with both congruence and uncertainty. A cell with value 1 or 0 has 100% congruence in models, predicting respectively a presence and an absence, while 0.5 represents cells where half of projections predict a presence, while the other half predict an absence. True skill statistics (TSS) and area under the receiver operating characteristic (ROC) values were calculated for each model, as well as a summary statistic for each of them. Models with both TSS and ROC values greater than the mean plus one standard deviation were kept to build the ensembles. This approach was pursued to evade subjective threshold adoptions.

We further explored the trends of climatic suitability values across past, current and future scenarios for each group of records (previously known, new and those from Mendoza). We extracted the suitability values for each presence record and calculated the average suitability in each group of records for each scenario. Values from the **PMN** dataset were used, as well as an average from the values considering all datasets separately ((**P** + **PN** + **PM** + **PMN**)/4). We compared both approaches and fitted a linear regression to make trends explicit.

### Niche equivalency and niche similarity tests

Ensemble models were a proxy of niche similarity and niche equivalency tests using the *ecospat* R package (Warren et al., 2008; Broennimann et al., 2012; Broennimann et al., 2018). Thus, we calculated for each pair of datasets the Schoener’s *D* statistics for niche overlap (Schoener, 1968) and a Hellinger distance-based metric (I) proposed by Warren et al., (2008). Both metrics range from 0 (no overlap) to 1 (indistinguishable) and are used to calculate similarity and equivalency p-values with simulated niches. The similarity test consists of trying to predict the niche of one dataset using the model generated for other dataset (Peterson et al., 1999) and has the assumption that niche conservatism is expected as a consequence of phylogenetic relationships and a finite rate of evolutionary divergence (Warren, 2008), thus, p-values lower than 0.05 represent niches that are more similar than expected by chance. This test returns two p-values for each pair of species based on the ensemble model built, e.g., for **P** will be used to predict **PN**, and ensemble model built for **PN** will be used to predict **P**. The equivalency test has an opposite view, where the assumption is that niches need to be indistinguishable to be equivalent, thus p-values greater than 0.95 represent niches that are more equivalent than expected by chance (Graham et al., 2004). This is reached through a random reallocation of occurrences of both datasets among their ranges. This routine was made through 100 simulated model niches (Dagnino et al., 2017), and for each simulated model niche we calculated the D niche overlap metric. We also explored a climatic space using the point localities values from the PCA-axes to discuss this topic.

## Results

### New distribution records and morphological features

In total, 27 distribution records were confirmed as *Opuntia bonaerensis* based on herbarium specimens. Of these, 19 are previously known from Argentina, mostly from the Buenos Aires province (15), one is from La Pampa and one is from Entre Ríos provinces; and two of them are from Mendoza, previously considered as from a non-natural distribution (discussed below). Eight records are newly generated from our field work and encompass the first records for the region, three being from south Brazil (Rio Grande do Sul state) and five from Uruguay (Colónia and Montevideo departments). All records are part of the Pampa or Río de La Plata grassland ecoregion, except those from Mendoza that are in the Puneña, Altoandean and Monte ecoregions. The full details regarding the distribution records reported here are presented in Fig. 1 and Table S2.

None of the unidentified materials previous deposited in the examined herbaria corresponded to *O. bonaerensis*, while three materials (UFP 24722, UFP 24723 and MPUC 13373) had misidentification as *O. bonaerensis* and were correctly assigned during our study as *O. elata* Salm-Dyck. Herbaria and field studies revealed morphological characters, such as bud flower apices, stigma color and inner pericarpel tissue color, as consistently useful to recognize *O. bonaerensis* (Fig. 4). However, spine production, although useful, was found to be a plastic character across populations.

### Phylogenetic analysis

Our alignment was 5,496 base pairs sequence in length with 51 parsimony-informative characters, 5,399 constant characters and 30 sites with gaps. The maximum likelihood inference depicted a tree with all nodes resolved on total bootstrap support (bs = 100) except for the position of *Opuntia rioplatense* Font as sister to the clade that includes *O. elata* + *O. megapotamica* Speg. (Fig. 2, bs = 52). The South American species of series *Elatae* formed a well-supported clade including all species proposed by Font (2014) except for *O. penicilligera* Speg., which was nested in the North American clade (Fig. 2). The three samples of *Opuntia bonaerensis*, including the records from Uruguay, Brazil and the specimen from the type locality in Argentina, formed a well-supported clade (bs = 100) sister to a clade with *O. elata*, *O. megapotamica* and *O. rioplatense*.

**Figure 2.**
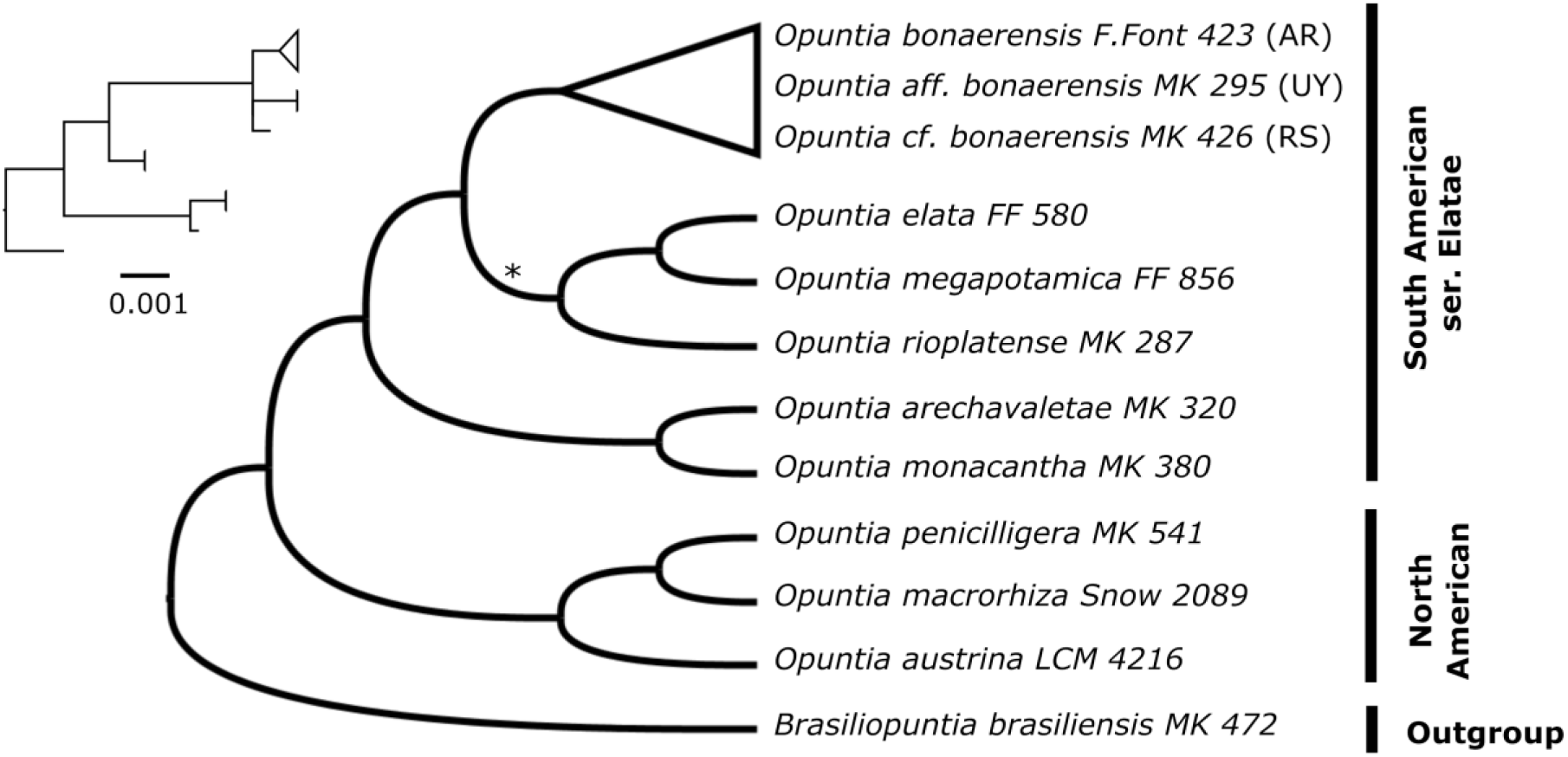
Maximum likelihood phylogenetic tree from RAxML analysis transformed in cladogram with the phylogram represented in small size with substitution rate scaled. All nodes have total bootstrap values (bs = 100) with exception to the asterisk denoting low bootstrap support (bs = 52). *Opuntia bonaerensis* from Argentina (*F.Font 423*) forms a well-supported clade with the two new records from Uruguay and Brazil (*MK 426* and *MK 295*).

### Ecological niche modelling

The models and projections for each climatic scenario and dataset generated are shown in Fig. 3. Statistics associated with ROC, TSS and number of models are summarized in Table 1. The dataset of the previous known distribution (**P**), excluding Mendoza records, projected a LGM (21 kya) distribution centered in Uruguay and adjacent areas as Entre Ríos province (Argentina) and Rio Grande do Sul (Brazil), occupying the continental shelf in the coast of Uruguay, shifting to central-east Argentina (especially the Buenos Aires province) in the MID (Middle Holocene (6 kya)). This distribution persisted until the current scenario, and future scenarios were predicted to keep suitability levels in the southern half of the distribution in the Buenos Aires province (RCP 2.6) and in the southernmost quarter of the distribution in the worst-case scenario (RCP 8.5). The previous known distribution plus the new records dataset (**PN**) projected a similar result for the LGM, but MID presented whole west-Uruguay as a suitable area, as well as connected areas in the southern Brazilian region of Pampean grasslands (Rio Grande do Sul). This pattern persists in the current scenario, where suitability of the models is divided between areas of previously known distribution and areas of Pampean region from Uruguay to Rio Grande do Sul, with an overall moderate suitability. Future scenario suitability shifts mostly to the coast in contrast to the **P** dataset, which presents a higher suitability in the inland. Despite the differences, both sets (**P** and **PN**) present a higher suitability along the coast of Argentina in the worst-case scenario (RCP 8.5). The third dataset (**PM**) shows a slightly different result, revealing the influence of the Mendoza records. Interior areas close to the piedmont of the Andes are more suitable than in other datasets. In the LGM, what is today called the Cuyo region in Argentina, had a greater suitability when comparing to aforementioned datasets. This effect persists through scenarios, increasing in RCP 2.6 and RCP 8.5, with lower suitability in southern Brazil. The last dataset (**PMN**), comprises a different perspective, where the species is predicted to have a wider distribution in LGM, occupying a connected area ranging from southern Paraguay, northeast and pampean Argentina, occurring throughout Uruguay, reaching parts of Brazil and the continental shelf. Within the MID scenario, the climate in Paraguay was less suitable, southern Brazil became widely climatically suitable and an overall southern shift started reaching the San Matías Gulf (Argentina). The current scenario in the **PMN** dataset suggested range contraction towards the province of Buenos Aires, with relicts in Uruguay and southern Brazil. Future projections show a tendency towards a southern and coastal shift, accompanied by an increase in suitability in the piedmont of the Andes. Generally, projections of all models unveiled a north-south distribution shift, ranging from a wider pattern in LGM to a more restricted distribution in an extreme climate change scenario (RCP 8.5).

**Figure 3.**
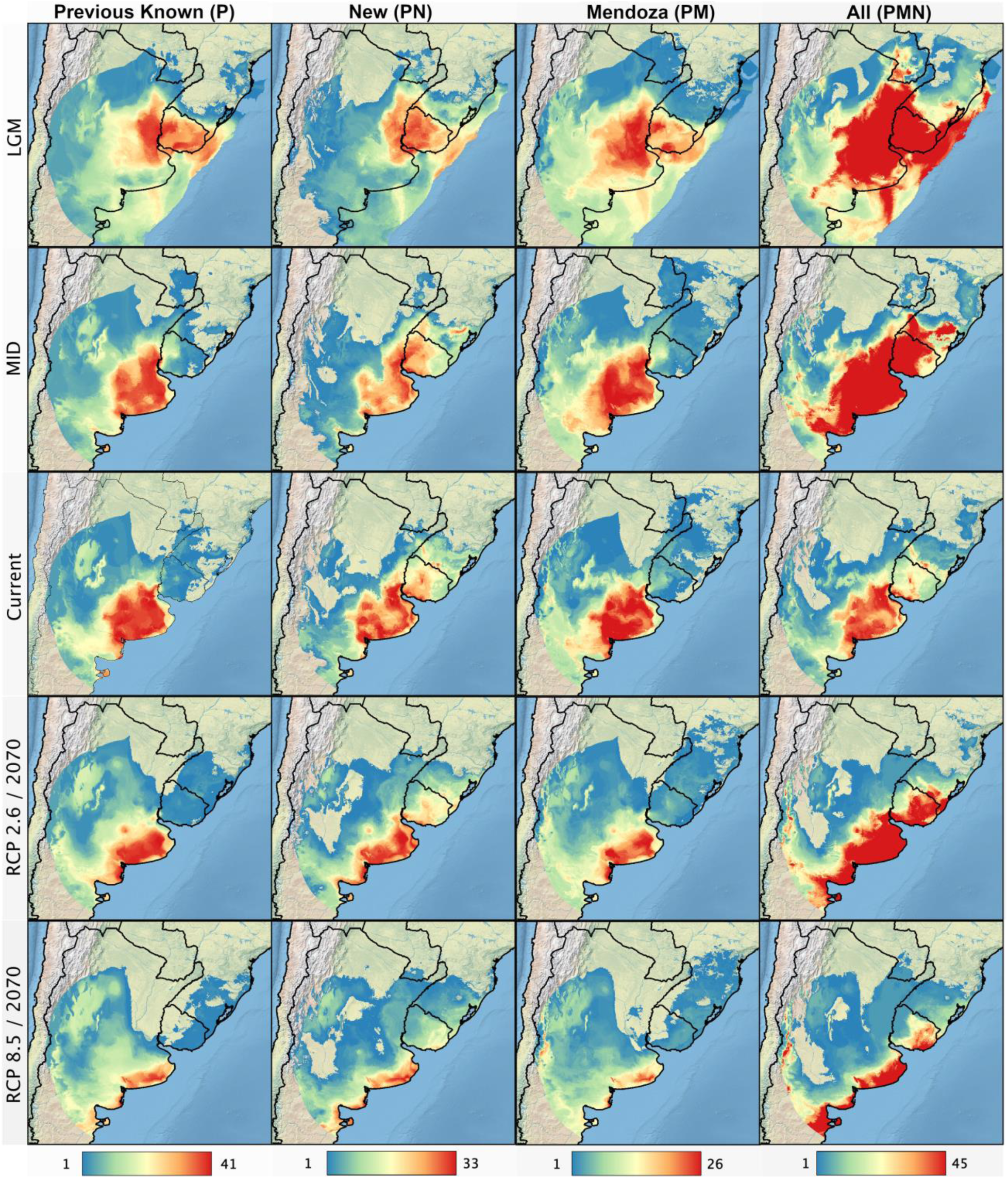
Projections for the *Opuntia bonaerensis* distribution to multiple scenarios (rows) considering different datasets (columns) using ensemble ecological niche models. LGM (Last Glacial Maximum, 21 ka); MID (Mid-Holocene, 6 ka); RCP (Representative Concentration Pathway) of 2.6 and 8.0 for future projections in 2070. Color scale represent agreement between models’ projections (cold colors represent low agreement; hot colors represent high agreement). (For interpretation of the references to color in this figure legend, the reader is referred to the web version of this article).

**Table 1.**
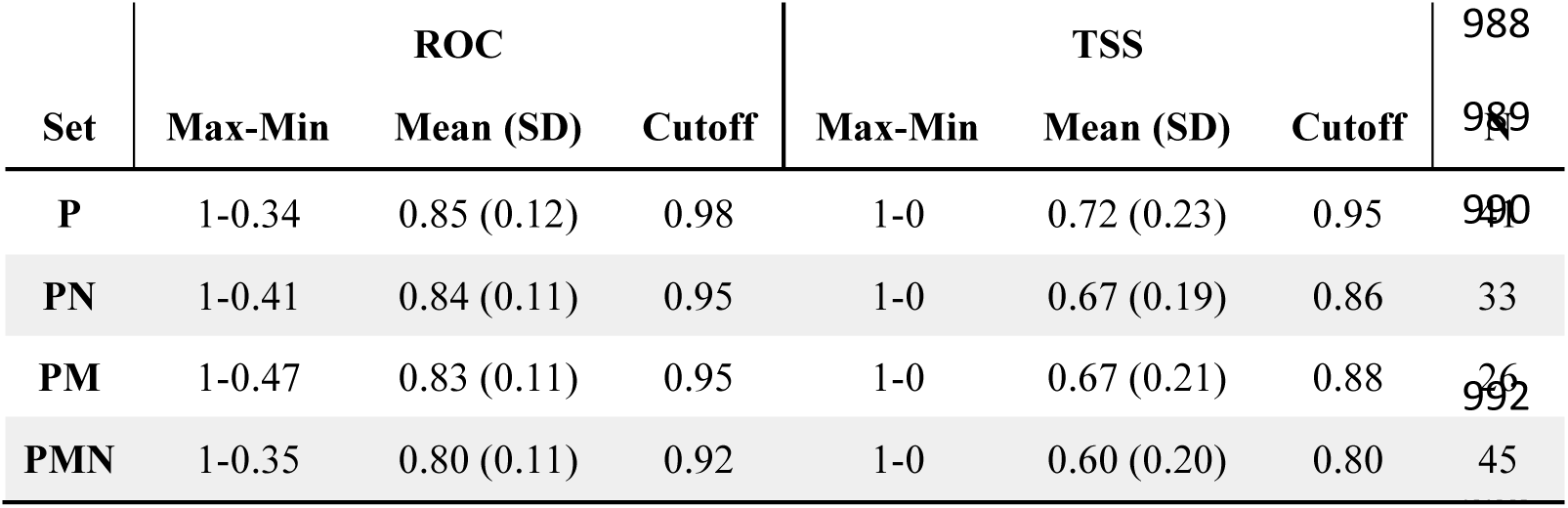
Ecological niche modelling results. Receiver Operating characteristic (ROC) and True Skill Statiscs (TSS) values. Cutoff values represent thresholds used to keep models in each dataset, summing a total of N models for each set.

Projections unveiled an uneven response to climatic change across populations (Fig. S1). Previously known records (Figure 1, green circles) had a higher mean climatic suitability in past and current scenarios, decaying in future scenarios, with values ranging from 0.55 and 0.8 in the LGM and MID, reaching a maximum in our current scenario, 0.93; decaying to 0.51 and 0.19 in RCP-2.6 and RCP-8.5. Likewise, new records (Figure 1, blue circles) had a higher mean climatic suitability in past and current scenarios, decaying in future scenarios, with values ranging from 0.49 and 0.7 in LGM and MID, reaching a maximum in current scenarios, 0.54 (average from all datasets) and 0.9 (**PMN** dataset); decaying to 0.37-0.58 and 0.17-0.23 in RCP-2.6 and RCP-8.5. On the contrary, records from Mendoza had a lower mean climatic suitability in past scenarios, while increasing in current and future scenarios, with values ranging from 0.11 and 0.17-0.3 in LGM and MID, reaching a maximum in current scenario, 0.4-0.6, with a little decay to 0.3-0.4 and 0.2 in RCP-2.6 and RCP-8.5 (Fig. S1).

### Niche equivalency and niche similarity

Niche equivalency tests returned higher p-values when comparing the **P** dataset with **PN** (p > 0.97) and **PMN** datasets (p > 0.99), which suggests that those sets have equivalent niches (Tab. 2). Contrarily, when comparing the **PM** dataset to every other set, as well as comparing the **PMN** set with the **PN** set, niche is less similar than expected by chance (Tab. 2). Climatic space showing the point localities from the PCA-axes also revealed that the Mendoza records add substantially different climatic information (Fig. S2). Despite that, niche similarity tests detected a small difference between sets, but with no statistical difference in any of them (Tab. 2).

**Table 2.**
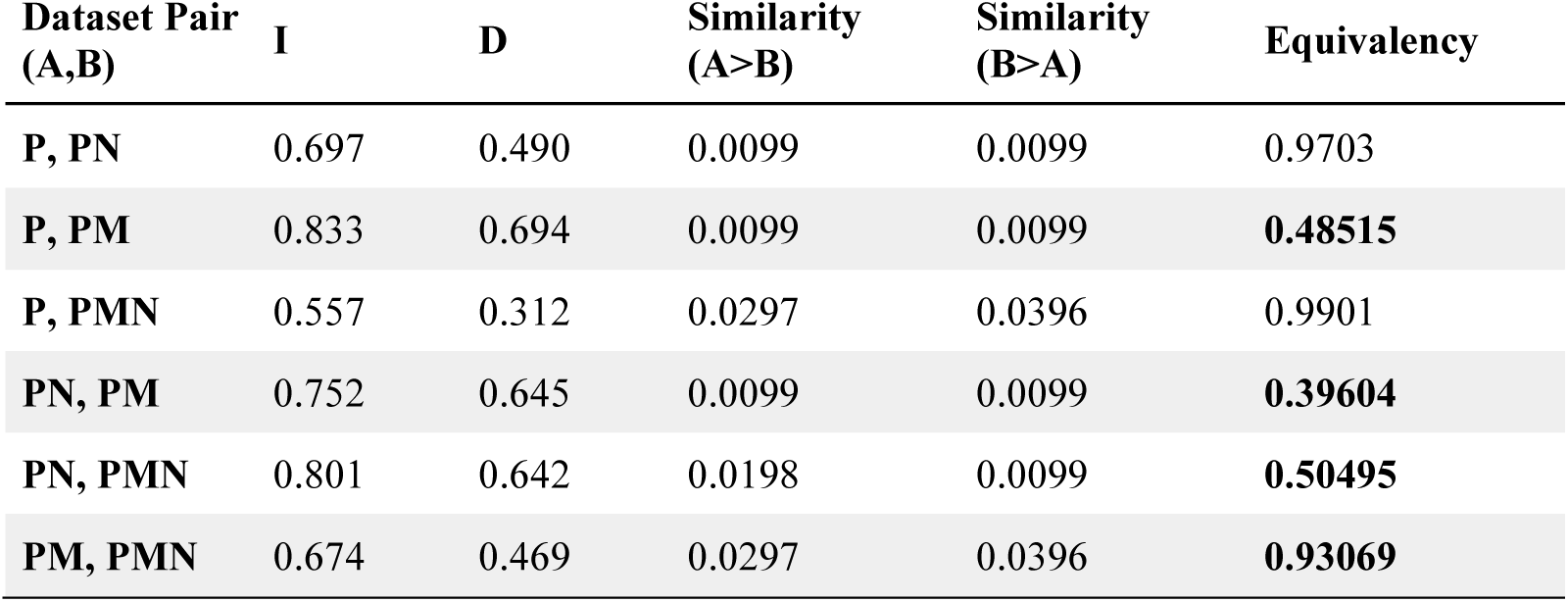
Niche similarity and equivalence tests. Values in Similarity and Equivalency columns are p-values. Bold equivalency values are those with p-values lower than 0.95, thus indicating that niches are less equivalent than expected by chance.

## Discussion

### Morphology and phylogenetic relationship of *Opuntia bonaerensis*

Species delimitation within *Opuntia* is well-known to be problematic (Rebman and Pinkava, 2001; Hunt, 2002; Powell et al., 2004; Majure et al., 2017). Several issues have contributed to this as 1) high amounts of hybridization resulting in mosaics of morphological features expressed by hybrid progeny, 2) morphologically variable species, where morphological characters are oftentimes dependent upon environmental variables (Griffiths, 1906; Reyes-Agüero et al., 2007) 3) poor specimen preparation and the lack of general collecting of species throughout their ranges, 4) the lack of basic biological data (e.g. chromosome counts, phenology, pollinators and floral biology, geographical distribution), 5) the absence of detailed studies regarding morphology across the distribution of species, and 6) the deficiency of phylogenetic data (Majure and Puente 2014). However, various efforts have been recently carried out to increase the knowledge of the group across all the range of distribution yielding valuable information to decisions regarding species delimitation (Powell et al., 2004; Majure et al., 2012a, 2012b, 2017; Font 2014; Realini et al., 2014a, 2014b; Las Peñas et al., 2017; Köhler et al., 2018; Martínez-González et al., 2019; Majure et al., 2020). Here, we highlighted the use of multiple approaches, such as molecular data, morphological characters, herbarium and field studies across geographical distribution as powerful tools to reveal accurate species identification in a problematic group.

*Opuntia bonaerensis* is easily recognized by an array of morphological features, such as the bright dark-green spathulate to long-ellipsoid stem segments, the acute flower buds with inner orange tepals and green stigma, and the short to long obconic fruits with the purple-wine inner pericarp tissue (Font 2014; Las Peñas et al., 2017) (Fig. 4), whereas the morphologically similar *O. elata* has oblong stem segments, creamy-white stigma lobes and pyriform fruits with green inner pericarpel tissue. The original description of *O. bonaerensis* suggested it to be a spineless or rarely 1-2 spine/areole-armed plant. Our revision of living plants and herbarium specimens revealed this to be a putatively plastic character, as some morphotypes were observed to be growing erect to curved, dark-reddish developing spines (Fig 4B). This feature has also been observed in some spineless morphotypes when grown under cultivation. Our extensive field work and herbaria examination suggested a transition of spineless morphotypes from southern Argentina to more spiny specimens in the newly reported populations. It is known that the spine color, as well the spine production, are in many instances phenotypically variable in *Opuntia* and change through time but can also be extremely diagnostic at the species level (Pinkava, 2003; Powell et al., 2004; Majure et al., 2017). Further studies should be carried out to determine if spine production in *O. bonaerensis* could be related to ecological factors leading to its development.

**Figure 4.**
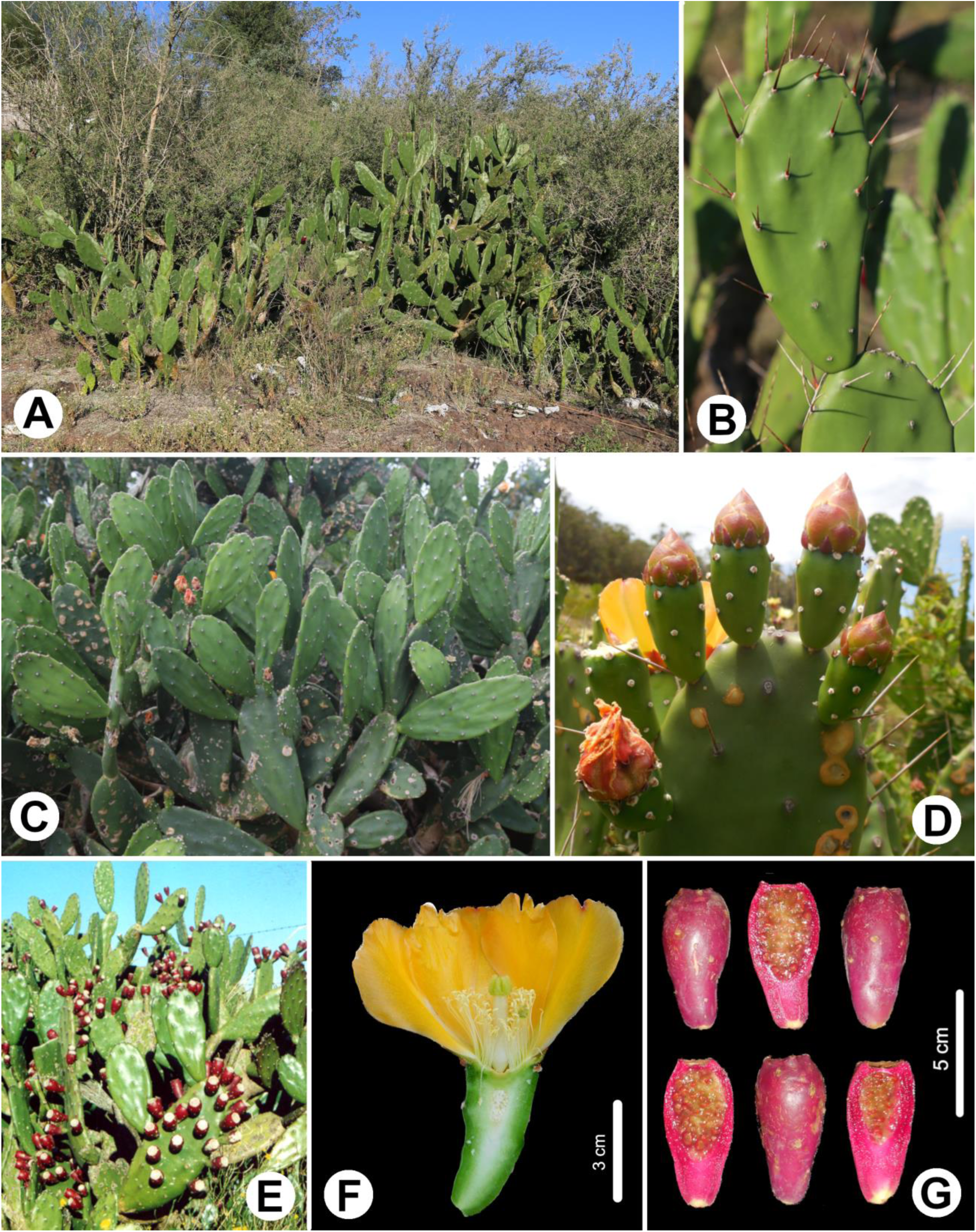
Morphological aspects of *Opuntia bonaerensis.* **A.** Typical habitat of the species occurring on rocky outcrops of the Pampa shrubby-grasslands (*M. Köhler 426*) **B.** Details of the cladodes with dark purple-reddish new spines when present in prickly morphotypes (*M. Köhler 424*) **C.** Typical spineless morphotype with elliptic to elongated-spatulate cladodes (*M. Köhler 295*) **D.** Acute flower buds with the dark-reddish external tepals (*M. Köhler* 295) **E.** Specimen from the type locality in Argentina (*F. Font 423*) **F.** Flower in transverse section showing the green stigma and the obconic ovary (*M. Köhler* 295) **G.** Long to shortly obconic ripe fruits in transverse section showing the purple pulp (*M. Köhler 424*). (All photos from M. Köhler, except **E** from F. Font) (For interpretation of the references to color in this figure, the reader is referred to the web version of this article).

*Opuntia bonaerensis* is a resurrected taxon proposed from an attentive revision of the southern South American species of the Elatae series based on analyses of protologues, field work, cultivation of specimens and examination of herbarium materials (Font 2014). This circumscription was supported in previous molecular studies (Realini et al., 2014b), and the molecular analysis presented here reinforces the taxon as a distinct lineage. The species is closely related to *Opuntia elata*, with which it was previously synonymized, and it is nested in the clade of the other “orange-flowered” southern South American *Opuntia* species of the Elatae series, such as *O. megapotamica* and *O. rioplatense* (Fig. 2). The morphological characters that have been now adopted to delimit these species (e.g., bud flower apices, stigma and inner pericarpel tissue color) appear to be strongly informative and reflect those evolutionary lineages.

The inclusion of all species proposed by Font (2014) as part of *Opuntia* series *Elatae* Britton and Rose (= ser. *Armatae* K. Schum.) in our phylogenetic analysis revealed that the series is not appropriately circumscribed. Font (2014) included seven species in the series and made a tentative inclusion of *Opuntia penicilligera* Speg. to accommodate a taxon that has been historically controversial and with minor affinities to the rest of the southern South American *Opuntia*. Previously treated in the *Sulphureae* series (Britton and Rose 1919), Las Peñas et al., (2017) maintained it in the series *Elatae*, but as suggested in our analysis, recent studies have shown that the species is phylogenetically nested in the North American *Humifusa* clade, and is likely derived from the *O. macrorhiza* species complex (Majure et al. 2020).

### An ancient distribution and now relictual in Brazil and Uruguay

Exploring the **P** dataset, ensembled models and projections to the past (LGM) clearly indicate that the potential distribution of *Opuntia bonaerensis* is largely congruent with our new records of the species in Uruguayan and Brazilian pampean region, which was previously unexpected from herbarium and literature reviews. Combining the previous known distribution with the new records (**PN** dataset), the ensembled models and projections maintain the suitability of the past distribution (LGM) in these regions with minimal differentiation, confirmed by the niche equivalent and niche similarity tests showing that newly discovered populations occupy similar environmental niches, compared to Buenos Aires populations (Tab. 2). This allowed us to suggest that the extant populations in these regions can be from a putative Pleistocene distribution that is relictual in the present.

During the climatic fluctuations of the Quaternary, the Southern Hemisphere was not affected by extensive glaciations as in the Northern Hemisphere, but the impacts of the Pleistocene have been increasingly documented in the South American flora and revealed as an important epoch for diversification both at the generic, as well as at the species level (Rull, 2008; Ramos-Fregonezi et al., 2015; Turchetto-Zolet et al., 2013; Ramírez-Barahona and Eguiarte, 2013). In cacti, several studies uncovered Pleistocene events related with climatic oscillations and glacial/interglacial cycles as a decisive driver for disjunct distributions, microrefugia and diversification across disparate clades emphasizing Mesoamerica, as well central, eastern and southeastern South American regions, primarily in the Atlantic forest and *Cerrado* biomes (Ornelas and Rodríguez-Gomez, 2015; Bonatelli et al., 2014; Franco et al., 2017a, 2017b; Silva et al., 2018a). However, diversification and distribution patterns in southern South American regions during the Pleistocene have been neglected, especially in the Pampas and Chaco regions. Just recently, studies revealed the impacts of climatic oscillations (e.g. glacial/interglacial cycles, sea level changes) as a driver of speciation and distribution in Solanaceae and Passifloraceae grassland species of the Pampa and Chaco domains (Mäder et al., 2013; Fregonezi et al., 2013; Ramos-Fregonezi et al., 2015; Moreno et al., 2018; Giudicelli et al., 2019).

Here, we demonstrate for the first time that events of the Pleistocene may have also impacted the distribution of cacti in southern South America. The Chaco-Pampa Plain is the southern part of the vast South American deposition trough. The present topography of the region was formed through the last regression of the Miocene Paranaense Sea and in a great part of the Chaco-Pampa Plain Quaternary loess and loessoid deposits cover Pliocene fluvial sand (Kruck et al., 2011). Lithostratigraphical and paleoenvironmental interpretations based on fossils have suggested that during the 28 – 16 kya, an arid climate with very weak humid spells dominated the region favoring a xerophytic vegetation (Zarate and Fasano 1989; Barreda et al., 2007; Quattrocchio et al., 2008; Kruck et al., 2011). This is congruent with other evidences that aridity has played an important role in cactus diversification and distribution (Hershkovitz and Zimmer 1997; Ritz et al., 2007; Arakaki et al., 2011). Besides that, aridity in the Chaco-Pampa Plain was accompanied by a lowering mean temperature (Zarate and Fasano 1989; Quattrocchio et al., 2008), which suggests that cold tolerance is an important feature for plants surviving in these environments. This feature can be easily related with extant populations of *Opuntia bonaerensis,* which have a remarkable presence in southern Buenos Aires province where low temperatures are dominant, and in the subtropical grasslands of Uruguay and Rio Grande do Sul.

In Uruguay and Rio Grande do Sul (Brazil), some metamorphic and granitic formations were temporarily isolated during the Pleistocene and Holocene by marine ingressions, affecting population dynamics and leading to the diversification of plant species in these regions (Mäder et al., 2013; Longo et al., 2014; Moreno et al., 2018). The higher parts of the orographic system of these regions are speculated to have been refugia during population expansions and retractions in the interglacial/glacial cycles and marine ingressions (Rambo, 1954), and could also had acted as orogenetic barriers for population containment. The putative participation of extinct large-size mammals (megafauna) on long-distance dispersal via migrating herbivores (Janzen, 1986) should not be neglected, since seeds of *Opuntia* have been found in wooly mammoth (*Mammuthus*) dung, and megafauna fossils have been richly recovered along many rivers of Pampean region (Davis et al., 1984; Scanferla et al., 2012).

There are no fossils to aid in an absolute dating of the Cactaceae. Taxon sampling with representative fossils in outgroups has yielded estimates for the crown age of cacti to be around 28.6 (26.7–30.5) million years ago (Mya) (Arakaki et al., 2011), 26.88 (16.67–37.10) Mya (Hernández-Hernández et al., 2014), 28.8 (15.08–48.15) Mya (Magallón et al., 2015) and 42.5 (54.5–26.5) Mya (Silva et al., 2018a). Although these can be understood as of moderate age, the subsequent divergence and diversification in the family was generated by significant radiations occurring more recently throughout the mid to late Miocene, into the Mid-Pliocene and more recently in the Pleistocene (Ritz et al., 2007; Arakaki et al., 2011, Hernández-Hernández et al., 2014, Bonatelli et al., 2014; Majure et al., 2019). In *Opuntia*, previous studies proposed that the clade started to diverge 5.6 Mya (+/-1.9) in the Late Miocene, but all major extant clades diverged during the Pliocene with subsequent diversification and speciation fully nested into the Pleistocene (Majure et al., 2012a), which is largely congruent with our hypothesis of Pleistocene impacts in the geographical distribution of *O. bonaerensis* across southern South America areas. However, further studies should be carried out sampling more individuals per population and a larger set of molecular markers, including from the nuclear genome, to access a more precise history of the Pleistocene influences in cactus genetic differentiation and putative linkage to specific bioregions.

The records from Mendoza are not absolutely resolved yet. The region is not represented by the Pampean domain, but rather from an Andean-Patagonic domain characterized by the Puneña and Altoandean floras (Oyarzabal et al., 2018). These records were previously reported by Font (2014) with accurate species identification, who suggested that this seemingly anomalous distribution could be from non-natural dispersion, as minor ornamental uses of the species are known in home-gardens in the capital of the province. However, our new records, combined with the previously known distribution, revealed a congruent distributional pattern that includes mountainous areas of southern Brazil, Uruguay and Argentina - known as the Neotropical Peripampasic Orogenic Arc – that has been reported for an array of animal taxa, such as spiders, scorpions, harvestmen and moths (Ferretti et al., 2012; Silva et al., 2018b), suggesting new insight for the historical biogeography of *Opuntia bonaerensis* that must be further explored.

### Implications for conservation

Brazil, Uruguay and Argentina are signatories of the Convention on Biological Diversity (CBD), following the Global Strategy for Plant Conservation (GSPC) and are directly dealing with plant knowledge, use and conservancy in their territories (Sharrock et al., 2018). Enormous efforts have been undertaken in Brazil to achieve some targets for the development of a functional and widely accessible list of all known plant species of the country (Forzza et al., 2010). However, as the country has long been acknowledged as a world leader in floristic diversity, it is clear that many gaps in our knowledge of the flora still need to be filled (Mittermeier and Mittermeier, 1997; Forzza et al., 2012; BFG, 2015).

*Opuntia* is a representative genus that exemplifies the increasing of local knowledge regarding its biodiversity. In the first attempt of an authoritative census of the Brazilian flora with scientific credibility to guide conservation planning, just one native species of *Opuntia* was reported as occurring in the country (Forzza et al., 2010; Forzza et al., 2012). Later, increasing the efforts to field studies, collections and preparation of materials for herbaria, more than ten species have been documented (Carneiro et al., 2016; Köhler et al., 2018; Zappi and Taylor, 2019; Köhler et al., *in prep.*).

Here, we provide the first report and confirm the presence of *Opuntia bonaeren*sis for the Brazilian and Uruguayan floras. With these new data, we augment the known distribution of the species, previously treated as endemic to Argentina, and expand the conservation efforts for the species. Rio Grande do Sul state has its own Red List of endangered flora, which helps to protect species from the different threats that plants and specially cacti suffer, and is frequently updated (Rio Grande do Sul, 2014). Although *O. bonaerensis* is one of the dozens of *Opuntia* species that have not been yet officially evaluated for its conservation status on the IUCN Red List, its assessment for the local Red List of Rio Grande do Sul is highly recommended for the next revision of the List, considering the limited populations and the ecological significance of the species in the region.

*Opuntia bonaerensis* is an endemic species of the Pampa biome and the Río de la Plata grassland, one of the largest continuous grassland ecoregions in the Americas covering the vast plains of central-eastern Argentina, Uruguay and part of southern Brazil (Andrade et al., 2018). This is a diverse and historically neglected region for conservation, with less than 3% of its territory under protection, that only now is receiving increasing efforts and strategies for its conservation (Krapovickas and Giacomo, 1998; Overbeck et al., 2015; Oliveira et al., 2017; Andrade et al., 2018; Vieira et al., 2019). In general, our projections for future scenarios under climate change revealed a north-south distribution shift when comparing to past and current models, with accumulation in the extreme-southern portion of the modern records (Fig. 3). These regions are poorly covered by protected areas, considering only the Natural Reserve of Bahía San Blas and the Ernesto Tornquist Provincial Park (both in Buenos Aires, ARG) as important protected areas that would contain *O. bonaerensis* distributions in worst-case projections.

The records from Mendoza, which are from non-cultivated populations, are in a distinct climatic condition, confirmed by our niche equivalence and similarity tests. It is important to note that there are only two records of difference between the **PM** and **P** datasets, which may bias the results of similarity tests. A proper way to do that should be comparing sets with no shared records, which is not possible in this case due to the small number of records in Mendoza. Curiously, ensemble models using the **P** and **PN** datasets revealed a slightly suitability of presences under the future (RCP 2.6 and RCP 8.5) and current scenarios, respectively, for regions encompassing the Mendoza province (Fig. 3). This, plus the trend of increasing suitability averages for Mendoza records (Fig. S1), suggests that the Mendoza region may act as a future refuge for *O. bonaerensis* under climate change scenarios.

### Conclusions

This study confirms for the first time the presence of *Opuntia bonaerensis* for the Brazilian and Uruguayan flora using molecular phylogenetics and morphology, extending the known distribution of the species and expanding the conservation efforts and strategies for it. Our update on the distribution of *O. bonaerensis* is coincident with the Neotropical Peripampasic Orogenic Arc and suggests new insights for the historical biogeography of the species that must be further explored. The assembled ecological niche models using four different datasets of presence records suggested that the newly revealed Brazilian and Uruguayan populations are putative relicts of a Pleistocene distribution, illuminating for the first time that climatic oscillations during the last 21,000 years may have played an important role in cactus distribution in southern South American Pampean-Chaco regions. Our analyses of climatic suitability trends revealed that the region of Mendoza, previously assumed to be from a non-natural distribution, may act as a future refuge for *O. bonaerensis* under the climate change scenarios explored. Further phylogeographic approaches, sampling more individuals per population and populational genetic markers should be pursued to reveal a detailed history of the species through its now better-known distribution.

### Author contributions

**M.K**: Conceptualization, Methodology, Formal analyses, Investigation, Data Curation, Writing - Original Draft, Writing - Review & Editing, Funding acquisition, Project administration; **L.F.E**.: Methodology, Formal analyses, Writing - Original Draft, Writing - Review & Editing; **F.F.**: Investigation, Data Curation, Writing - Review & Editing; **T.T.S.C.**: Writing - Review & Editing; **L.C.M.**: Methodology, Writing - Review & Editing, Supervision, Funding acquisition.

## Acknowledgments

MK thanks to Cassio R. da Silva, Josimar Külkamp, Marcos V.B. Soares, Anderson S. Mello and Philipy Weber for providing help during fieldwork; Ethiéne Guerra for providing comments during submission; Alexandra Elbakyan for help providing access to references; and Lucas Kaminski for valuable discussions during the manuscript writing. MK is grateful to the American Society of Plant Taxonomists (ASPT), Cactus and Succulent Society of America (CSSA), International Association for Plant Taxonomy (IAPT) and IDEA WILD for supporting part of the research here reported. MK also thanks the Brazilian National Council for Scientific and Technological Development (Conselho Nacional de Desenvolvimento Científico e Tecnológico - CNPq) for his PhD scholarship, and the PDSE/CAPES for support his period as Visiting Researcher at the Florida Museum of Natural History (FLMNH, UF, USA). This study was also financed in part by the Coordenação de Aperfeiçoamento de Pessoal de Nível Superior – Brasil (CAPES) - Finance Code 001 and start-up funds to LCM from Florida Museum of Natural History. Two anonymous reviewers provided comments on the manuscript to improve it.

---

**Figure S1.**
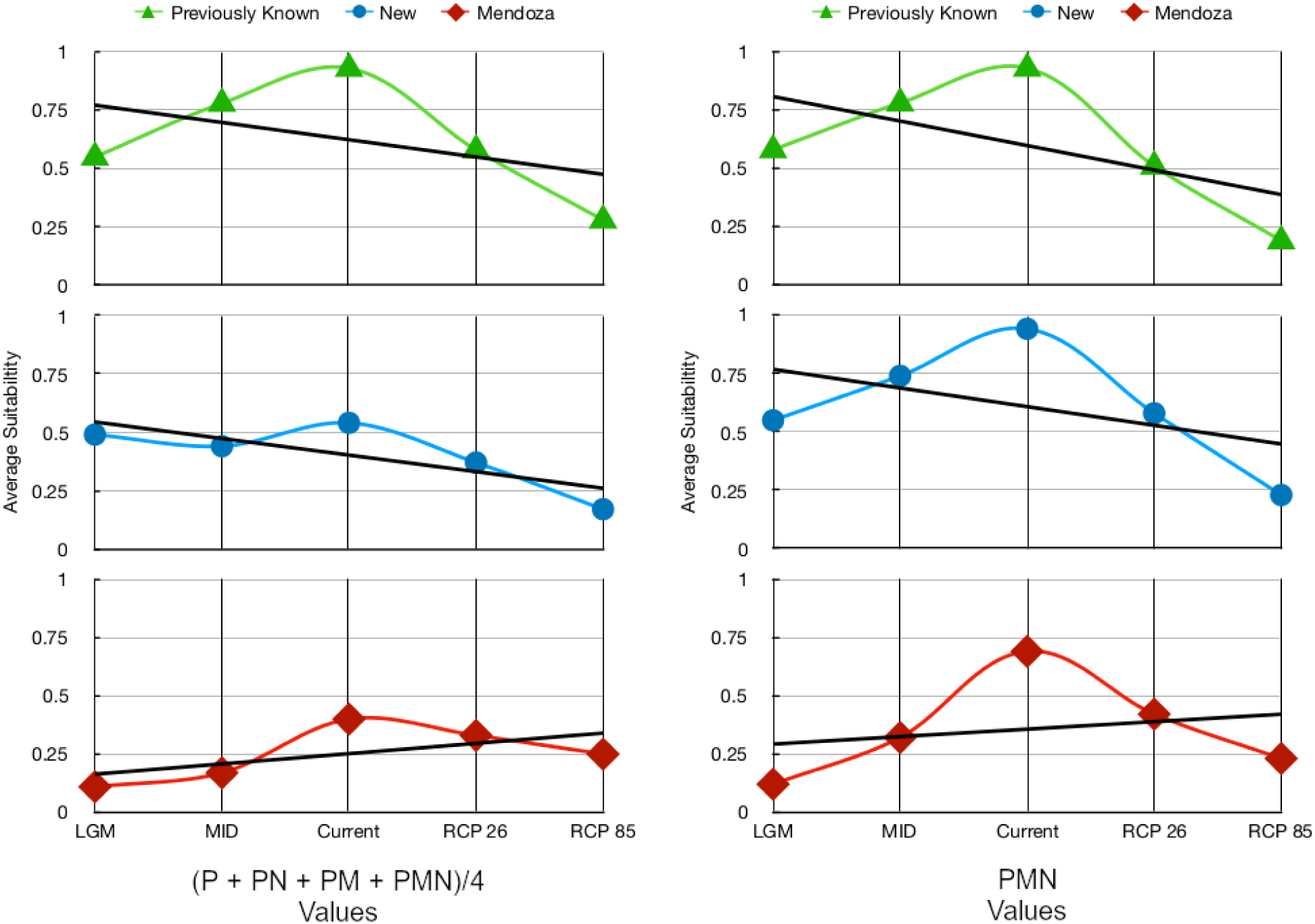
Trends of climatic suitability of presence records across different projected scenarios. Left graph represented average values from all datasets, while right graph represented average values from the **PMN** dataset. Both approaches revealed a trend of climatic increase suitability for Mendoza records and a decrease for the previously known and new records.

**Figure S2.**
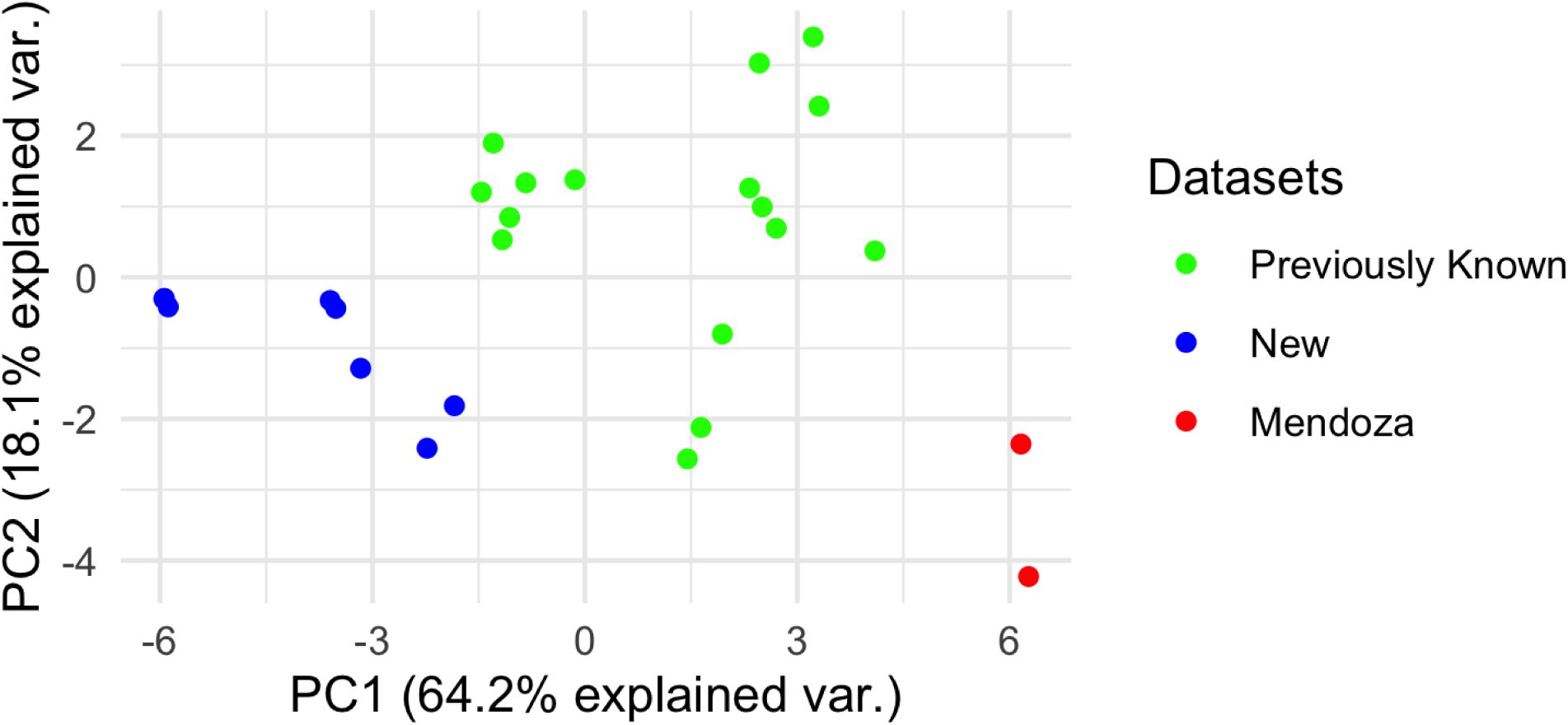
Climatic space graph from the PCA-axes showing that Mendoza records added distinct climatic features for ecological niche modelling.

## Table captions

**Table S1.**
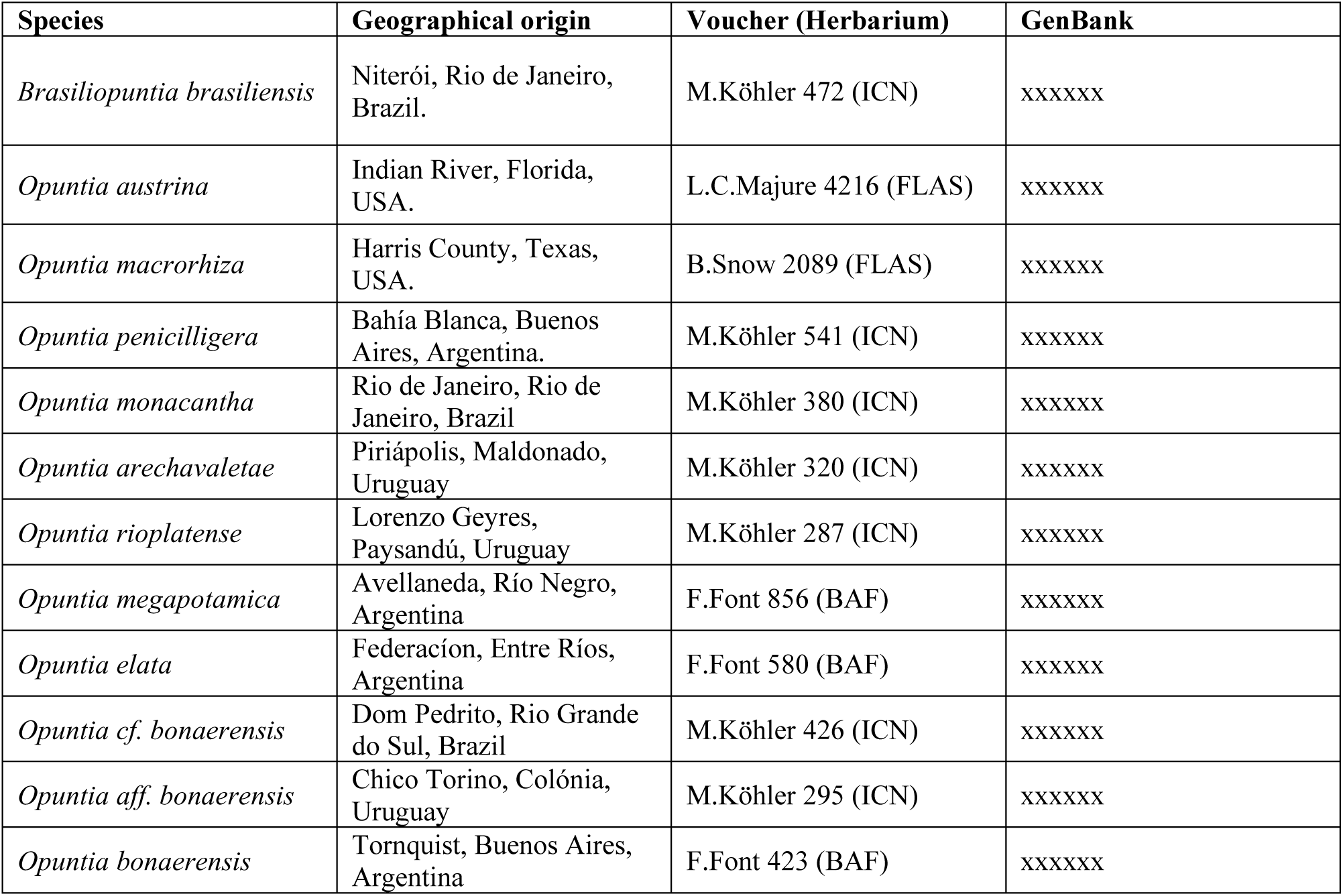
Sampled material for molecular analyses.

**Table S2.** Distribution records used for Ecological Niche Models (ENM).

| **Dataset** | **Geographical origin** | **Geographical coordinates** | **Voucher (Herbarium)** |
| --- | --- | --- | --- |
| Previously known | Chivilcoy, Buenos Aires, Argentina | -34.951167<br>-60.018778 | G.Logarzo, 556 (BAF, BCWL) |
| Previously known | Suipacha, Buenos Aires, Argentina | -34.669528<br>-59.744194 | F.Font 424 (BAF) |
| Previously known | Tandil, Buenos Aires, Argentina | -37.453000<br>-59.176278 | F.Font 587 (BAF) |
| Previously known | Zárate, Buenos Aires, Argentina | -34.062472<br>-59.161306 | F.Font 586 (BAF) |
| Previously known | Conhelo, La Pampa, Argentina | -35.941000<br>-64.273028 | G.Logarzo 551 (BCWL) |
| Previously known | San Andrés de Giles, Buenos Aires, Argentina | -34.395972<br>-59.336389 | F.Font 422 (BAF) |
| Previously known | Tornquist, Buenos Aires, Argentina | -38.234694<br>-62.407278 | F.Font 423 (BAF) |
| Previously known | Villarino, Buenos Aires, Argentina | -38.895194<br>-63.162861 | F.Font 584 (BAF) |
| Previously known | Campana, Buenos Aires, Argentina | *-34.210062<br>-58.958603 | Castellanos s.n. (BA 17378) |
| Previously known | Trenque Lauquen, Buenos Aires, Argentina | -36.061861<br>-62.829361 | F. Font 585 (BAF) |
| Previously known | Campana, Buenos Aires, Argentina | -34.221028<br>-58.897917 | G.Logarzo 463 (BCWL) |
| Previously known | González Chávez, Buenos Aires, Argentina | -38.041639<br>-60.109167 | G.Logarzo 528 (BCWL) |
| Previously known | Gualeguaychú, Entre Ríos, Argentina | -33.491250<br>-58.799583 | F.Font 419 (BAF) |
| Previously known | Bahía Blanca, Buenos Aires, Argentina | -38.473750<br>-62.287611 | Villamil 8826 (BBB) |
| Previously known | Bahía Blanca, Buenos Aires, Argentina | -38.656472<br>-62.223250 | Villamil 9006 (BBB) |
| Previously known | Tandil, Buenos Aires, Argentina | *-37.797667<br>-59.036167 | Spegazzini s.n. (BA 63450) |
| Previously known | Capital, La Pampa, Argentina | -36.590889<br>-64.211167 | G.Logarzo 553 (BAF, BCWL) |
| Mendoza – Previously known | San Carlos, Mendoza, Argentina | -33.961889<br>-69.181472 | G.Logarzo 544 (BAF, BCWL) |
| Mendoza – Previously known | Tunuyán, Mendoza, Argentina | -33.609889<br>-69.181472 | G.Logarzo 546 (BAF, BCWL) |

\* Geographical coordinates were generated by our assumption based on label data.

|  |  |  |  |
| --- | --- | --- | --- |
| New records | Dom Pedrito, Rio Grande do Sul,<br>Brazil | -30.995908<br>-54.624319 | M.Köhler 203<br>(ICN) |
| New records | Chico Torni, Colónia, Uruguay | -34.402439<br>-57.259072 | M.Köhler 295<br>(ICN) |
| New records | Colónia Valdense, Colónia, Uruguay | -34.342414<br>-57.215431 | M.Köhler 297<br>(ICN) |
| New records | Rincón del Pino, San José, Uruguay | -34.502392<br>-56.835381 | M.Köhler 300<br>(ICN) |
| New records | Montevideo, Montevideo, Uruguay | -34.840969<br>-56.409758 | M.Köhler 307<br>(ICN) |
| New records | Montevideo, Montevideo, Uruguay | -34.887081<br>-56.260525 | M.Köhler 315<br>(ICN) |
| New records | Dom Pedrito, Rio Grande do Sul,<br>Brazil | -30.952639<br>-54.654103 | M.Köhler 424<br>(ICN) |
| New records | Dom Pedrito, Rio Grande do Sul,<br>Brazil | -30.962692<br>-54.660094 | M.Köhler 426<br>(ICN) |

